# Gene polymorphism associated with Angiotensinogen(M235T), Endothelial lipase (584C/T) and susceptibility to coronary artery disease: A meta-analysis

**DOI:** 10.1101/2020.04.24.059295

**Authors:** Hongyan Zhao, Shan Hu, Jidong Rong

## Abstract

**Objective:** To explore the association between the variant M235 locus of angiotensinogen (AGT) gene, 584C/T locus of Endothelial lipase (EL) gene, and coronary artery disease (CAD) by meta-analysis.

**Methods:** The case-control studies on the association between AGT/EL gene polymorphism and CAD were collected through searching PubMed, EMbase, Web of Science, CNKI and Wanfang database up to March 1, 2020. Stata 15.0 software was used for analysis.

**Results:** A total of 29 articles met the inclusion criteria. After analyzing, it was found that the M235T polymorphism of AGT gene was associated with the occurrence of CAD. In the allele model (T vs. M), OR=1.38 (P < 0.05). In other heredity, there was also statistically significant. Subgroup analysis indicated that except the heterozygous genetic model of the Chinese population, other genetic models of the Caucasian and Chinese population were also statistically significant. The 584C/T polymorphism of EL gene was associated with the occurrence of CAD, with OR=0.83 (P < 0.05) in the allele model (T vs. C) and OR=0.80 (P < 0.05) in the dominant gene model. Also, in the allele model of Caucasian subgroup, OR=0.83 (P < 0.05), while in Asian subgroup, there was no statistically significant genetic model.

**Conclusion:** AGT M235 and EL 584C/T polymorphisms are associated with CAD susceptibility. The genotype TT, TC or allele T of AGT M235T and genotype CC or allele C of EL 584C/T might be the genetic risk factors for the development of CAD.

## 1 Introduction

Coronary artery disease (CAD), the incidence of which is increasing year by year in both developed and developing countries, is the leading cause of death and seriously endangers human health [1, 2]. With the increase of age, the incidence of CAD increases gradually [3]. Common risk factors, such as dyslipidemia, hypertension and diabetes, are affected by both environmental and genetic factors [4]. Also, these traditional risk factors could not directly lead to CAD, and there might be other factors directly leading to atheromatous plaques formation are related to genetic susceptibility. Moreover, increasing studies have demonstrated that the renin-angiotensin system (RAS) plays a vital role in CAD [5].

Renin is mainly secreted by renal juxtaglomerular cells, and it can catalyze the conversion of plasma pro-angiotensin (AGT) into Angiotensin I. Also, Angiotensin I can be further converted into angiotensin II, III, IV, to play the role of vasoconstriction. It has been found that AGT M235T polymorphism is closely related to the occurrence of cardiovascular disease. M235T refers to the substitution of nucleotide T at location 704 of the 2nd exon by C, resulting in the conversion of amino acid at position 235 from methionine (M) to threonine (T). Some studies have confirmed that Angiotensinogen M235T polymorphism is closely related to the severity of coronary artery disease [6]. Van et al. [7] have suggested that the haplotype of angiotensinogen gene is associated with coronary heart disease in familial hypercholesterolemia. While, Angiotensinogen M235T polymorphism is not associated with coronary artery disease in diabetic patients [8]. Khatami et al. [9] suggested that allele T of the AGT gene would increases the risk of CAD, while Zhu et al. [9] indicated that allele T would not increase the risk of CAD.

Related studies have shown that low plasma high-density lipoprotein cholesterol (HDL-C) concentration is closely related to the incidence of coronary heart disease (CAD). In contrast, high HDL-C levels can reduce the risk of CAD [11]. The concentration of HDL-C in plasma is not only affected by environmental factors, but also highly hereditary. Current genetic studies indicate that the genetic heterogeneity of HDL-C is associated with the gene polymorphism of endothelial lipase (LIPG), which could regulate the metabolism of HDL [12]. Endothelial lipase (EL), a member of the triglyceride lipase family, has high homology with Lipoprotein lipase (LPL) and Hepatic lipase (HL), and plays a critical role in the regulation of lipid metabolism, especially in the process of HDL-C metabolism. The main activity of endothelial lipase is phospholipase activity, and HDL granules are its preferred substrate. Plasma endothelial lipase activity affects human high-density lipoprotein metabolism and coronary artery risk factors [13]. The relationship between EL 584C/T polymorphism and CAD susceptibility has been discussed, but there is not consistent conclusion. Solim et al. [14] believed that allele T of EL gene was a protective factor of CAD, but Xie et al. [15] did not find this association in the Chinese population.

Furthermore, to compare different research results more scientifically and objectively, meta-analysis is used to conduct a comprehensive study on this issue. On this basis, a meta-analysis of the genotypic data from all eligible surveys in recent years was administered to more accurately evaluate the relationship between Angiotensinogen (M235T), Endothelial (584C/T) polymorphism and susceptibility to coronary artery disease, thus providing evidence-based medicine for cardiovascular clinic.

## 2 Methods

### 2.1 Literature search

A combination of following term, Angiotensinogen, Endothelial lipase, myocardial infarction, coronary artery disease, acute myocardial infarction, cardiovascular disease, ischemic heart disease, coronary heart disease, cute coronary syndrome and polymorphism, were used to systematically search the relevant literatures on PubMed, EMbase, Web of Science, China National Knowledge Infrastructure (CNKI) and Wanfang databases up to March 1, 2020. The research language was unlimited.

### 2.2 Inclusion and exclusion criteria

#### 2.2.1 Inclusion criteria

1) case-control study, 2) CAD was defined as coronary heart disease, coronary artery disease, myocardial infarction, acute Myocardial infarction, acute coronary syndrome, cardiovascular disease, 3) the number of genotypes should be provided in both the case group and the control group.

#### 2.2.2 Exclusion criteria

1) No specific amount of genotypes was provided, or the data of each genotype cannot be obtained by calculation, 2) repeated studies of the same ethnic group, 3) NOS score was less than 6.

### 2.3 Data extraction and methodological quality

According to the Newcastle-Ottawa Scale (NOS) [16], the full text was carefully read and evaluated. The literature below six stars was of low quality, and the research above six stars was of high quality, among which only the research above 6 stars was included. The evaluation was conducted independently by two evaluators according to the uniform quality standard, and they extracted the data from the literature and then cross-checked it. When they encountered some differences, they solve these problems through discussion, or with the help of a third party. The extracted data included the number of AGT M235T and EL 584C/T genotypes in the case group and the control group, first author, publication time, source of the control group, country.

### 2.4 Statistical analysis

The odds ratio (OR) and its 95% confidence interval were used to evaluate the association between AGT/EL gene polymorphism and CAD. The data were merged and analyzed by statistical software Stata 15.0. Q test was applied to verify the heterogeneity of the included studies. When I^2^ ≥ 50% or P ≤ 0.05, heterogeneity was considered to exist, the random effect model (REM) was selected; when I^2^ < 50% and *P* > 0.05, heterogeneity was deemed to be low, and the fixed effect model (FEM) was used for data integration. Z test was used to test the significance of the combined OR value. The funnel plot was performed to evaluate publication bias, and the criterion was whether the funnel plot was symmetric or not. If the funnel chart was asymmetric, there might be publication bias. Egger’s Test was also used for checking publication bias. Subgroup analysis was carried out to assess whether Hardy-Weinberg equilibrium (HWE) was satisfied in the Ethnicity and control groups. Finally, the sensitivity of the results was analyzed to determine the robustness of the research results.

## 3 Results

### 3.1 Search results

According to the inclusion and exclusion criteria, 29 articles were collected in this meta-analysis, including 16 studies on the relationship between AGT M235T polymorphism and CAD susceptibility [17-32], and 13 studies on EL 584C/T polymorphism and CAD susceptibility [33-45]. The specific literature screening process was displayed in figure 1. The characteristics of the study, the distribution frequency of genotypes and the results of the literature quality score were shown in Table 1 and Table 2.

**Table. 1.** The basic characteristics of AGT M235T polymorphism included in the literature

| First author | year | Country | End point | source of controls | Cases |  |  | Controls |  |  | HWE (control) | NOS score |
| --- | --- | --- | --- | --- | --- | --- | --- | --- | --- | --- | --- | --- |
|  |  |  |  |  | TT | MT | MM | TT | MT | MM |  |  |
| Ko YL | 1997 | China | CAD | HB | 225 | 36 | 6 | 279 | 54 | 4 | 0.454 | 8 |
| Chen D | 1998 | China | MI | PB | 40 | 13 | 4 | 32 | 31 | 13 | 0.258 | 8 |
| Sheu WH | 1998 | China | CAD | PB | 75 | 26 | 1 | 107 | 37 | 1 | 0.247 | 7 |
| Xie Y | 2001 | China | CAD | HB | 69 | 29 | 8 | 45 | 30 | 11 | 0.109 | 7 |
| Zhu TB | 2002 | China | CAD | HB | 56 | 48 | 14 | 54 | 42 | 10 | 0.661 | 8 |
| Gu WD | 2003 | China | CAD | PB | 86 | 31 | 12 | 53 | 30 | 7 | 0.568 | 8 |
| Zhu TN | 2004 | China | CAD | HB | 105 | 75 | 12 | 54 | 36 | 8 | 0.355 | 8 |
| Li L | 2004 | China | CAD | PB | 49 | 60 | 11 | 25 | 41 | 14 | 0.689 | 7 |
| Ren J | 2005 | China | CAD | HB | 67 | 24 | 9 | 31 | 26 | 13 | 0.087 | 7 |
| Yang JP | 2013 | China | CAD | HB | 73 | 27 | 10 | 34 | 29 | 17 | 0.031 | 7 |
| Huang YZ | 2013 | China | CAD | HB | 104 | 23 | 10 | 54 | 51 | 20 | 0.184 | 7 |
| Al-Hazzani | 2014 | Caucasian | CAD | HB | 73 | 98 | 54 | 31 | 50 | 29 | 0.342 | 8 |
| Bonfim a | 2016 | African-Brazilians | CAD | HB | 71 | 104 | 25 | 27 | 28 | 11 | 0.4234 | 8 |
| Bonfim b | 2016 | Caucasian-Brazilians | CAD | HB | 103 | 146 | 73 | 25 | 67 | 34 | 0.439 | 8 |
| Khatami | 2017 | Iran | CAD | HB | 15 | 50 | 83 | 10 | 31 | 94 | 0.004 | 8 |
| Isordia-Salas | 2018 | Mexico | MI | PB | 6 | 98 | 138 | 10 | 62 | 170 | 0.163 | 8 |
| Zhu M | 2019 | China | CAD | HB | 23 | 11 | 3 | 123 | 42 | 5 | 0.545 | 8 |

Note:HWE: Hardy-Weinberg equilibrium; CAD: coronary artery disease; MI:,myocardial infarction.HB:hospital based;PB:population based;NOS:Newcastle-Ottawa Scale

**Table. 2.** The basic characteristics of Endothelial lipase 584C/T polymorphism included in the literature

| First author | Year | country | End point | source of controls | CAD |  |  | Control |  |  | HWE (control) | NOS score |
| --- | --- | --- | --- | --- | --- | --- | --- | --- | --- | --- | --- | --- |
|  |  |  |  |  | CC | CT | TT | CC | CT | TT |  |  |
| Rimm | 1992 | America | CAD | PB | 129 | 117 | 16 | 240 | 239 | 40 | 0.060 | 8 |
| Colditz | 1997 | America | CAD | PB | 115 | 110 | 16 | 224 | 214 | 39 | 0.220 | 8 |
| Tjonneland | 2007 | Danmark | CAD | PB | 509 | 406 | 83 | 837 | 673 | 133 | 0.840 | 8 |
| Zhu JC | 2007 | China | CAD | PB | 186 | 56 | 0 | 150 | 46 | 0 | 0.063 | 7 |
| Tang NP | 2008 | China | CAD | HB | 174 | 85 | 6 | 125 | 122 | 18 | 0.100 | 8 |
| Zhang L | 2009 | China | CAD | HB | 493 | 379 | 69 | 405 | 333 | 56 | 0.264 | 7 |
| Cai GJ | 2014 | China | CAD | HB | 84 | 45 | 6 | 97 | 64 | 5 | 0.146 | 8 |
| Xie L | 2015 | China | CAD | HB | 160 | 116 | 11 | 216 | 139 | 12 | 0.065 | 8 |
| Elnaggar | 2018 | Egypt | CAD | HB | 58 | 23 | 2 | 17 | 21 | 4 | 0.492 | 8 |
| Solim | 2018 | Turkish | CAD | HB | 40 | 27 | 6 | 26 | 33 | 14 | 0.545 | 8 |
| Shimizu | 2007 | Japan | CAD | PB | 70 | 36 | 1 | 54 | 50 | 3 | 0.030 | 8 |
| Vergeer | 2010 | Netherlands | CAD | PB | 97 | 413 | 87 | 207 | 857 | 207 | 0.000 | 8 |
| Toosi | 2015 | Iran | CAD | HB | 67 | 70 | 3 | 28 | 46 | 6 | 0.029 | 8 |

Note:HWE: Hardy-Weinberg equilibrium; CAD: coronary artery disease; MI:,myocardial infarction.HB:hospital based;PB:population based;NOS:Newcastle-Ottawa Scale.

**Figure. 1.**
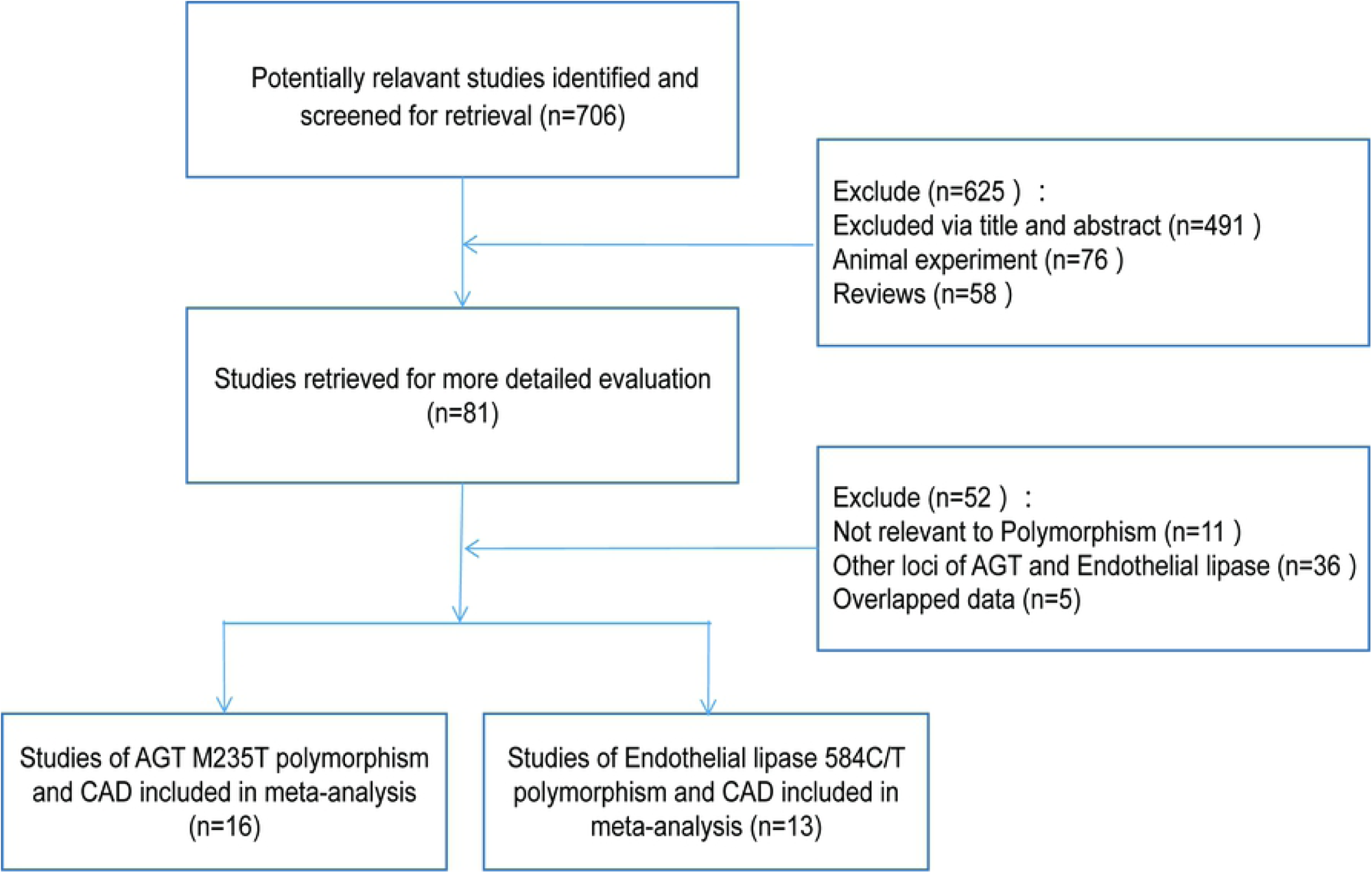
A flow diagram of the study selection process

### 3.2 Meta-analysis results

#### 3.2.1 Comparison of alleles

The main results of Meta-analysis of AGT M235T polymorphism and CAD susceptibility were shown in Table 3 and figure 2, while the main findings of Meta-analysis of EL 584C/T polymorphism and CAD susceptibility were shown in Table 4 and figure 3. The comparison of allele T and allele M at M235T locus of AGT gene suggested that there were significant differences in heterogeneity among different studies (I^2^ = 69.2%, P < 0.05). Therefore, the random effect model was applied. The results indicated that OR=1.38 (95%CI:1.15∼1.65), the difference was statistically significant. The same conclusion was obtained from the subgroup analysis of Chinese and Caucasian populations. The allele model was also statistically significant after the control group was removed from the study that did not meet HWE. The funnel plot (Fig. 4A) was symmetrical. Egger’s test results showed that P>0.05, suggesting no publication bias.

**Table. 3.** The Results of Meta-analysis of AGT M235T polymorphism and CAD susceptibility

| Gene model | Subgroup | n | OR | 95CI | p | I<sup>2</sup> | P for Heterogeneity | Model | P for Publication bias (Egger) |
| --- | --- | --- | --- | --- | --- | --- | --- | --- | --- |
| T vs. M | Overall | 17 | 1.38 | 1.15~1.65 | 0.001 | 69.2 | 0.000 | REM | 0.459 |
|  | China | 12 | 1.42 | 1.08~1.87 | 0.011 | 76.7 | 0.000 | REM | 0.767 |
|  | Caucasian | 4 | 1.37 | 1.16~1.61 | 0.000 | 0.0 | 0.601 | FEM | 0.532 |
|  | HWE satisfied | 15 | 1.32 | 1.09~1.60 | 0.005 | 68.9 | 0.000 | REM | 0.590 |
| TT+TM vs. MM | Overall | 17 | 1.51 | 1.28~1.80 | 0.000 | 17.3 | 0.251 | FEM | 0.764 |
|  | China | 12 | 1.53 | 1.15~2.02 | 0.003 | 32.8 | 0.128 | FEM | 0.205 |
|  | Caucasian | 4 | 1.52 | 1.21~1.90 | 0.000 | 0.0 | 0.402 | FEM | 0.337 |
|  | HWE satisfied | 15 | 1.43 | 1.19~1.73 | 0.000 | 16.0 | 0.274 | FEM | 0.670 |
| TT vs. TM+MM | Overall | 17 | 1.42 | 1.11~1.83 | 0.006 | 68.1 | 0.000 | REM | 0.884 |
|  | China | 12 | 1.54 | 1.12~2.12 | 0.008 | 73.3 | 0.000 | REM | 0.458 |
|  | Caucasian | 4 | 1.39 | 1.03~1.89 | 0.033 | 32.5 | 0.217 | FEM | 0.395 |
|  | HWE satisfied | 15 | 1.36 | 1.04~1.78 | 0.023 | 69.3 | 0.000 | REM | 0.891 |
| TT vs. MM | Overall | 17 | 1.56 | 1.15~2.12 | 0.004 | 43.9 | 0.027 | FEM | 0.759 |
|  | China | 12 | 1.66 | 1.05~2.12 | 0.029 | 54.8 | 0.011 | REM | 0.272 |
|  | Caucasian | 4 | 1.45 | 1.01~2.08 | 0.042 | 0.0 | 0.425 | FEM | 0.443 |
|  | HWE satisfied | 15 | 1.46 | 1.05~2.03 | 0.026 | 43.8 | 0.036 | REM | 0.746 |
| TM vs. MM | Overall | 17 | 1.32 | 1.10~1.59 | 0.003 | 2.5 | 0.425 | FEM | 0.037 |
|  | China | 12 | 1.07 | 0.79~1.46 | 0.650 | 0.0 | 0.755 | FEM | 0.201 |
|  | Caucasian | 4 | 1.48 | 1.16~1.88 | 0.001 | 50.7 | 0.107 | FEM | 0.334 |
|  | HWE satisfied | 15 | 1.25 | 1.03~1.53 | 0.027 | 4.3 | 0.425 | FEM | 0.055 |

Note:REM:random-effects model;FEM:fixed-effects model;OR:odds ratio;HWE:Hardy-Weinberg equilibrium.

**Table. 4.** The Results of Meta-analysis of Endothelial lipase 584C/T polymorphism and CAD susceptibility

| Gene model | Subgroup | n | OR | 95CI | p | I<sup>2</sup> | P for Heterogeneity | Model | P for Publication bias (Egger) |
| --- | --- | --- | --- | --- | --- | --- | --- | --- | --- |
| T vs. C | Overall | 13 | 0.83 | 0.73~0.95 | 0.006 | 69.5 | 0.000 | REM | 0.011 |
|  | Asian | 6 | 0.84 | 0.66~1.07 | 0.151 | 74.5 | 0.001 | REM | 0.462 |
|  | Caucasian | 7 | 0.83 | 0.70~0.98 | 0.029 | 68.9 | 0.004 | REM | 0.000 |
|  | HWE satisfied | 10 | 0.84 | 0.72~0.98 | 0.031 | 72.9 | 0 | REM | 0.064 |
| TT+TC vs. CC | Overall | 13 | 0.80 | 0.68~0.95 | 0.009 | 66.8 | 0.000 | REM | 0.013 |
|  | Asian | 6 | 0.80 | 0.60~1.06 | 0.116 | 73.5 | 0.002 | REM | 0.443 |
|  | Caucasian | 7 | 0.81 | 0.65~1.01 | 0.057 | 64.0 | 0.011 | REM | 0.003 |
|  | HWE satisfied | 10 | 0.82 | 0.68~0.99 | 0.036 | 70.1 | 0.000 | REM | 0.072 |
| TT vs. TC+CC | Overall | 12 | 0.87 | 0.75~1.02 | 0.078 | 32.3 | 0.132 | FEM | 0.100 |
|  | Asian | 5 | 0.92 | 0.68~1.23 | 0.563 | 43.8 | 0.130 | FEM | 0.668 |
|  | Caucasian | 7 | 0.86 | 0.72~1.02 | 0.090 | 32.8 | 0.177 | FEM | 0.009 |
|  | HWE satisfied | 9 | 0.90 | 0.75~1.08 | 0.244 | 37.2 | 0.121 | FEM | 0.173 |
| TT vs. CC | Overall | 12 | 0.73 | 0.55~0.98 | 0.034 | 52.9 | 0.016 | REM | 0.043 |
|  | Asian | 5 | 0.75 | 0.39~1.45 | 0.396 | 60.0 | 0.040 | REM | 0.647 |
|  | Caucasian | 7 | 0.70 | 0.49~1.00 | 0.050 | 54.9 | 0.038 | REM | 0.001 |
|  | HWE satisfied | 9 | 0.74 | 0.53~1.05 | 0.092 | 56.9 | 0.017 | REM | 0.141 |
| TC vs. CC | Overall | 13 | 0.84 | 0.72~0.97 | 0.019 | 57.2 | 0.005 | REM | 0.017 |
|  | Asian | 6 | 0.81 | 0.62~1.04 | 0.100 | 66.4 | 0.011 | REM | 0.429 |
|  | Caucasian | 7 | 0.87 | 0.72~1.05 | 0.143 | 50.7 | 0.058 | REM | 0.008 |
|  | HWE satisfied | 10 | 0.85 | 0.71~1.00 | 0.055 | 57.2 | 0.005 | REM | 0.083 |

Note:REM:random-effects model;FEM:fixed-effects model;OR:odds ratio;HWE:Hardy-Weinberg equilibrium.

**Figure. 2.**
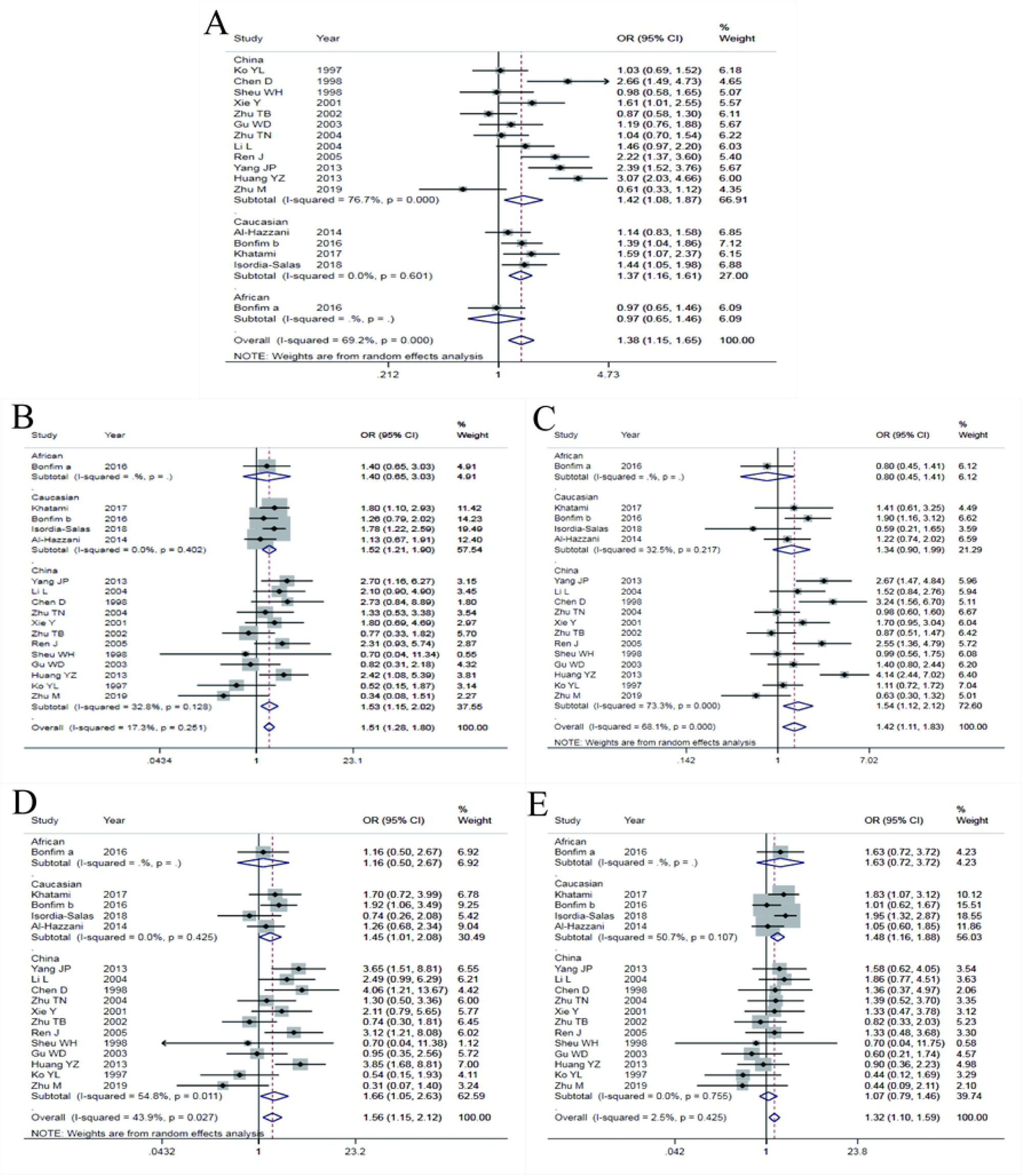
Forest plots of AGT M235T polymorphism and CAD susceptibility(A:Allele model; B:dominant model; C:recessive model; D:homozygous model; E:heterozygous model)

**Figure. 3.**
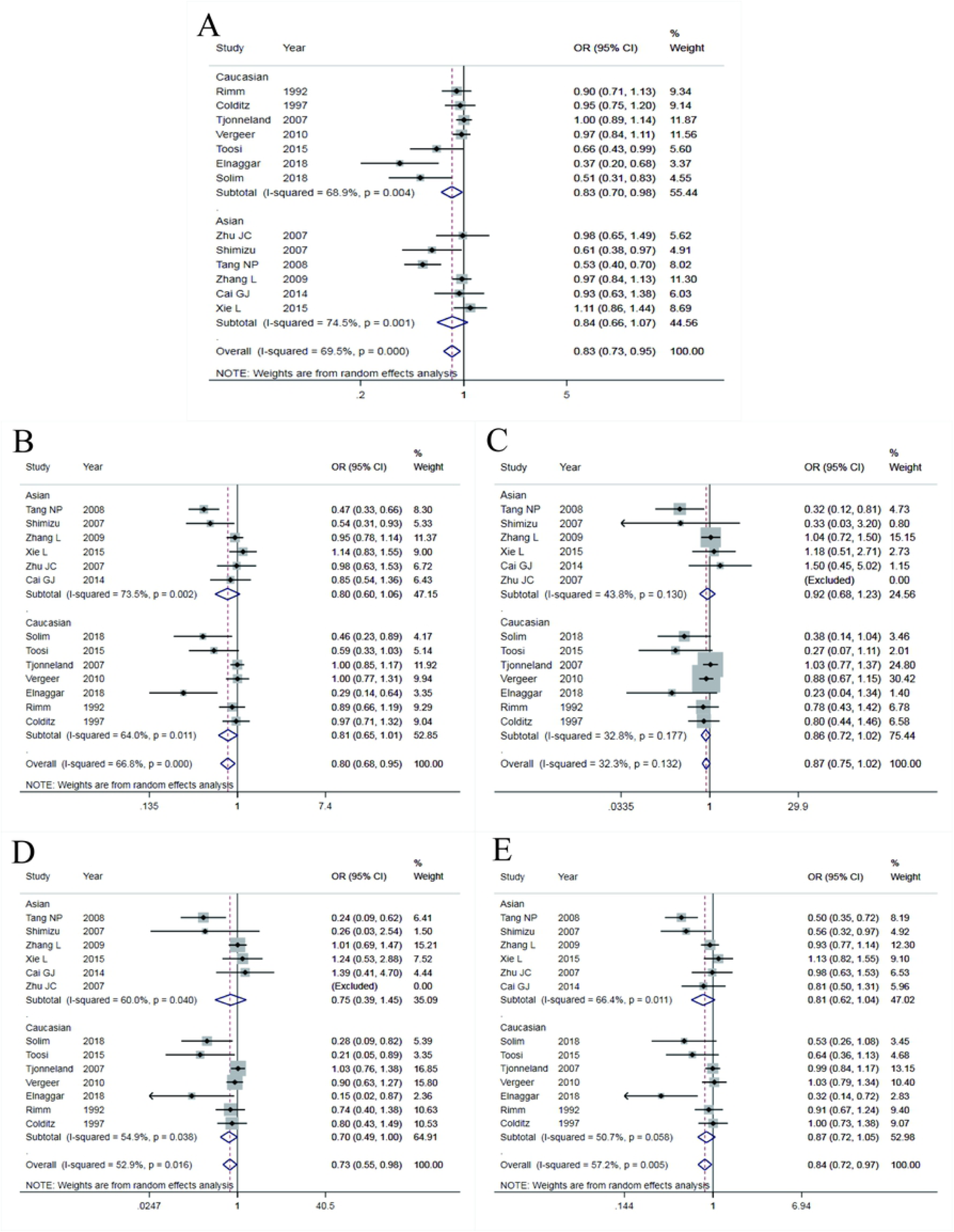
Forest plots of Endothelial lipase 584C/T polymorphism and CAD susceptibility(A:Allele model; B:dominant model; C:recessive model; D:homozygous model; E:heterozygous model)

**Figure. 4.**
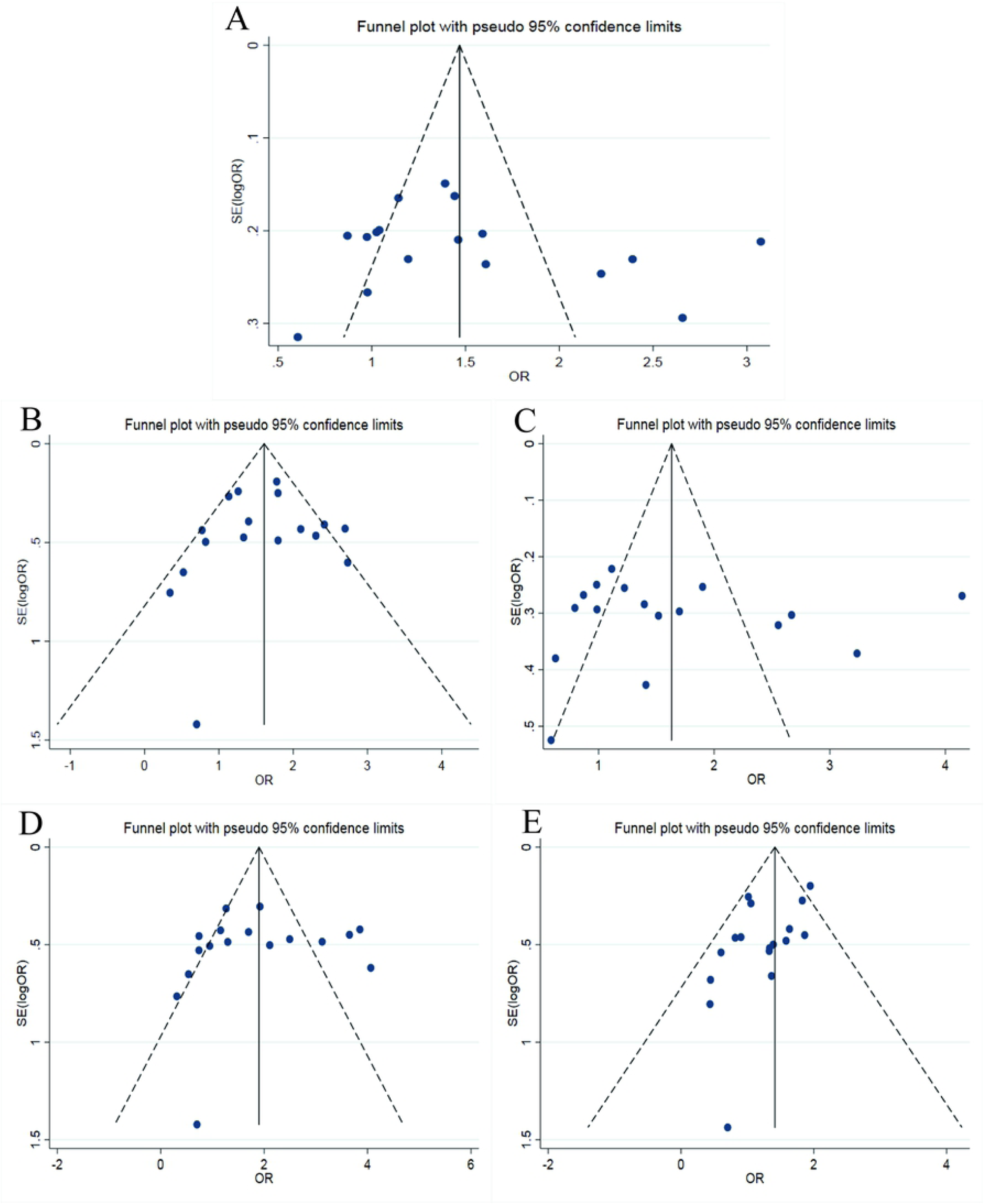
Funnel plots of AGT M235T polymorphism and CAD susceptibility(A:Allele model; B:dominant model; C:recessive model; D:homozygous model; E:heterozygous model)

Compared allele C with allele T at 584C/T locus of EL gene, the heterogeneity between studies was significant (I^2^=69. 5%, P< 0.05). Therefore, the random effect model was used for data merging. The results showed that OR=0.83 (95%CI:0.73∼0.95), the difference was statistically significant. The analysis of the Caucasian population subgroup and the subgroup without the control group that did not meet HWE showed statistically significant differences. In the Asian population, the allele model was not statistically significant. The funnel plot (see in figure.5A) was generally symmetrical, and the P-value of Egger’s test was less than 0.05, indicating a particular publication bias.

#### 3.2.2 Comparison of dominant gene inheritance models

The comparison of genotype TT+MT with genotype MM at M235T locus of the AGT gene showed that the heterogeneity between studies was not significant (I^2^ = 17.3%, P > 0.05). Therefore, the fixed effect model was used for data merging. The results showed that OR=1.51 (95%CI:1.28∼1.80), the difference was statistically significant. The same conclusion was obtained in the subgroup analysis of Chinese and Caucasian populations. After excluding the study that the control group did not meet HWE, the dominant gene model was also statistically significant. The funnel diagram was symmetrical (see in fig.4B). Egger’s test results indicated that there was no publication bias with P> 0.05.

The heterogeneity between the genotype TT+TC and genotype CC of EL 584C/T was significant (I^2^=66.8%, P < 0.05), so, the random effect model was used for data integration. The results showed that OR=0.80 (95%CI:0.68∼0.95), the difference was statistically significant. Excluding the subgroup analysis of the control group that did not meet HWE, the difference was statistically significant. In the subgroup analysis of the Caucasian and Asian populations, the dominant gene model was not statistically significant. The symmetry of the funnel plot (Fig.5B) was general, P-value of Egger’s test was less than 0.05, indicating that there was a particular publication bias.

#### 3.2.3 Comparison of recessive gene inheritance models

The comparison of genotype TT with genotype MT+MM at M235T locus of AGT gene showed that the heterogeneity between the studies was significant (I^2^ = 68.1%, P < 0.05), then, the random effect model was used to integrate data. The results indicated that OR=1.42 (95%CI:1.11∼1.83), the difference was statistically significant. The same conclusion was reached in the subgroup analysis of Chinese and Caucasian populations. The recessive gene model was also statistically significant after removing the control groups that did not satisfy HWE. The funnel plot was symmetrical, shown in fig.4C. Egger’s test results suggested that there was no publication bias (p > 0.05).

Compared genotype TT with genotype TC+CC at 584C/T locus of EL gene, there was no significant heterogeneity between studies (I^2^ = 32.3%, P > 0.05), and the fixed effect model was used. The results showed that there was no significant difference with OR=0.87 (95%CI:0.75∼1.02). Subgroup analysis that removed the control group that did not meet HWE showed no statistically significant difference. The recessive gene model was also not statistically significant in the Caucasian and Asian subgroups. The funnel plot (fig.5c) was symmetrical. The P-value of Egger’s Test was higher than 0.05, indicating a low publication bias.

#### 3.2.4 Comparison of homozygous genetic models

The comparison between genotype TT and genotype MM at M235T locus of the AGT gene showed no significant heterogeneity between studies (I^2^ = 43.9%, P > 0.05). Therefore, the fixed effect model was applied. The results showed that OR=1.56 (95%CI:1.15∼2.12), the difference was statistically significant. The same conclusion was obtained from the subgroup analysis of Chinese and Caucasian populations. After excluding the study that the control group did not meet HWE, the homozygous genetic model was statistically significant. The funnel plot was symmetrical (see in fig.4D). Egger’s test results suggested that there was no publication bias (p > 0.05).

Compared genotype CC with genotype TT at 584C/T locus of EL gene, there was significant heterogeneity (I^2^ = 52.9%, P < 0.05), then, the random effect model was applied. The results showed that OR=0.0.73 (95%CI:0.55∼0.98), the difference was statistically significant. Subgroup analysis that removed the control group that did not meet HWE showed no statistically significant difference. In the subgroup analysis of Caucasian and Asian populations, the heterogeneity was not significantly decreased, and the homozygous gene model was not statistically significant. The symmetry of the funnel chart shown in figure 5D was general, and the P-value of Egger’s Test was slightly less than 0.05, indicating partial publication bias.

**Figure. 5.**
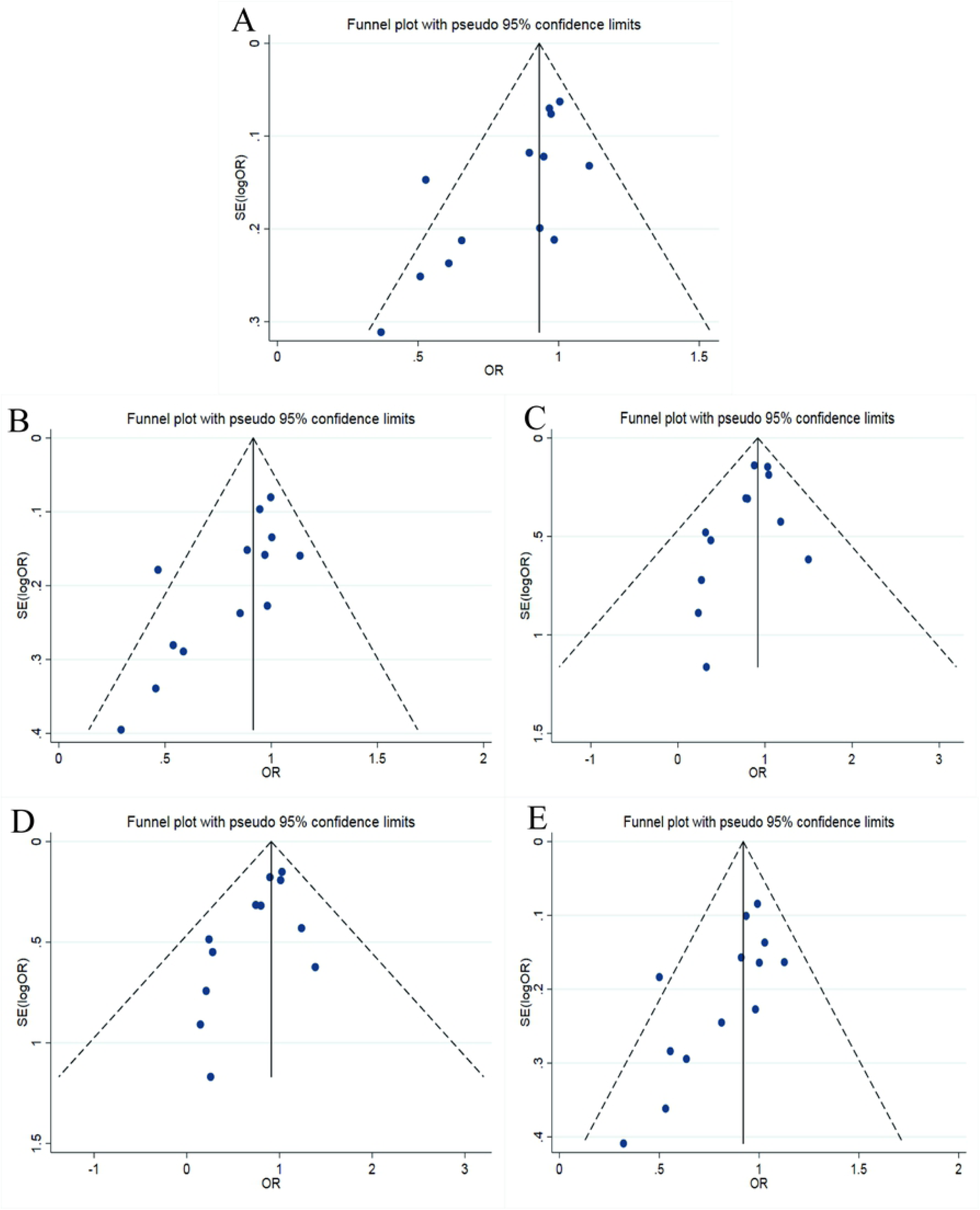
Funnel plots of Endothelial lipase 584C/T polymorphism and CAD susceptibility(A:Allele model; B:dominant model; C:recessive model; D:homozygous model; E:heterozygous model)

#### 3.2.5 Comparison of heterozygous genetic models

The comparison of genotype TM at M235T locus of the AGT gene with genotype MM suggested that the heterogeneity was not significant (I^2^ = 2.5%, P > 0.05). Therefore, the fixed effect model was used for data merging. The results showed that OR=1.32 (95%CI:1.10∼1.59), the difference was statistically significant. The same conclusion was obtained in the Caucasian population subgroup after removal of the control group that did not satisfy HWE. While, the heterozygous genetic model had no statistical significance in the Chinese population. The symmetry of the funnel plot was general (see in fig.4E). Egger’s test results showed that p<0.05, indicating that there was a partial publication bias.

Compared genotype CC with genotype TC at 584C/T locus of EL gene, significant heterogeneity was found (I^2^ = 57.2%, P < 0.05), then the random effect model was applied for data merging. The results showed that OR=0.84 (95%CI:0.72∼0.97), the difference was statistically significant. Subgroup analysis that removed the control group that did not meet HWE showed no statistically significant difference. In the subgroup analysis of the Caucasian and Asian populations, the heterozygous gene model was also not statistically significant. The funnel plot shown in Fig.5E was generally symmetrical. Egger’s test result indicated partial publication bias with P-value that was slightly less than 0.05,

### 3.3 Sensitivity analysis

The result of susceptibility analysis between AGT M235T polymorphism and susceptibility to CAD was shown in Fig.6. It indicated there was a statistically significant change in the heterozygous genetic model after elimination of one study. In contrast, no statistically significant change manifested in the allele model and other genetic models after removing the single literature, suggesting that the result was robust.

**Figure. 6.**
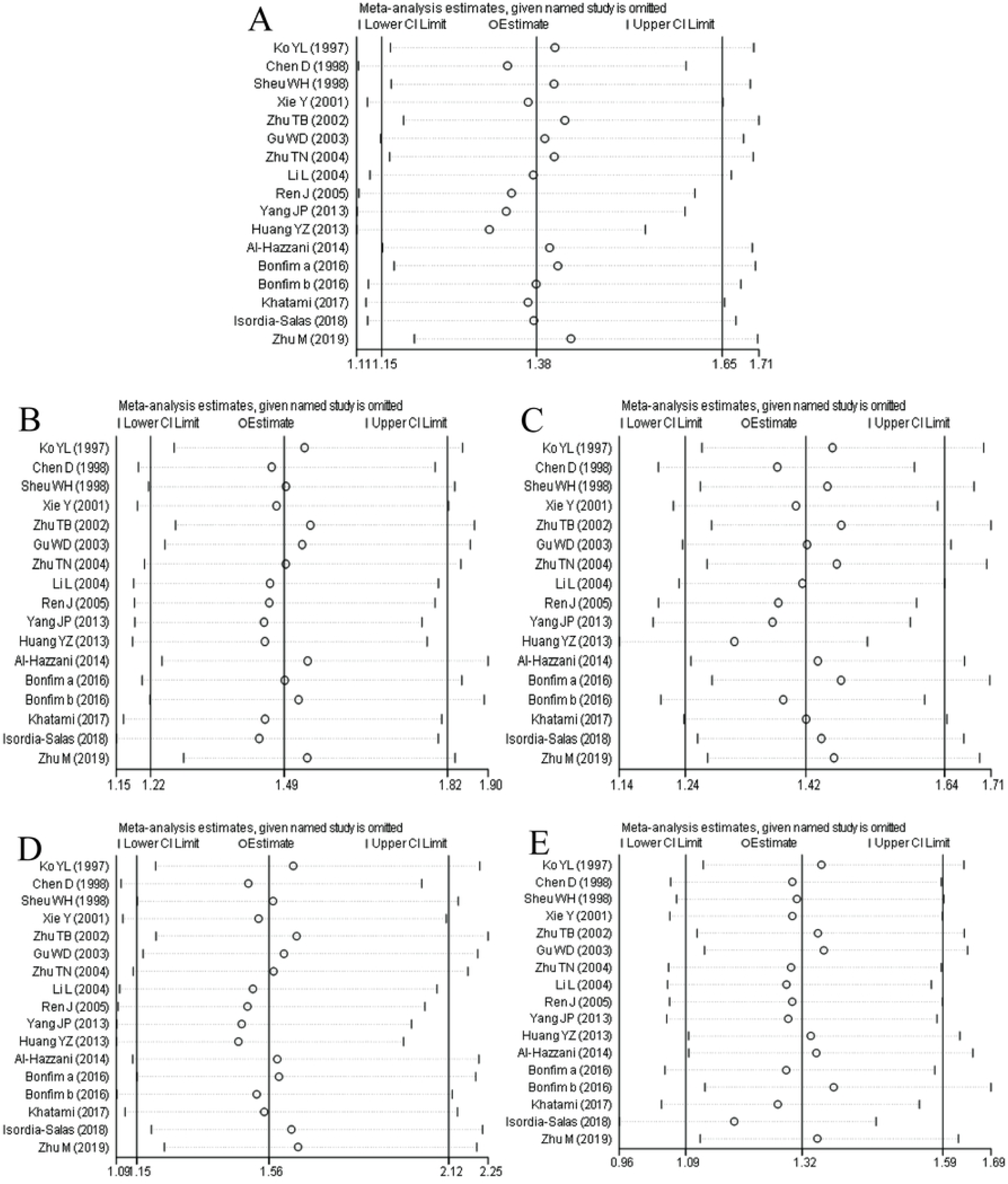
The result of susceptibility analysis between AGT M235T polymorphism and susceptibility to CAD(A:Allele model; B:dominant model; C:recessive model; D:homozygous model; E:heterozygous model)

The sensitivity analysis result of EL 584C/T polymorphism and CAD susceptibility was shown in Fig.7. It showed that there was no statistically significant change in the allele model after a single study was eliminated, indicating that the result of the allele model was robust. The dominant gene model, recessive gene model, homozygous gene model and heterozygous gene model had 1, 1, 4, 1 study, respectively, and they were excluded one by one; the differences were statistically significant.

**Figure. 7.**
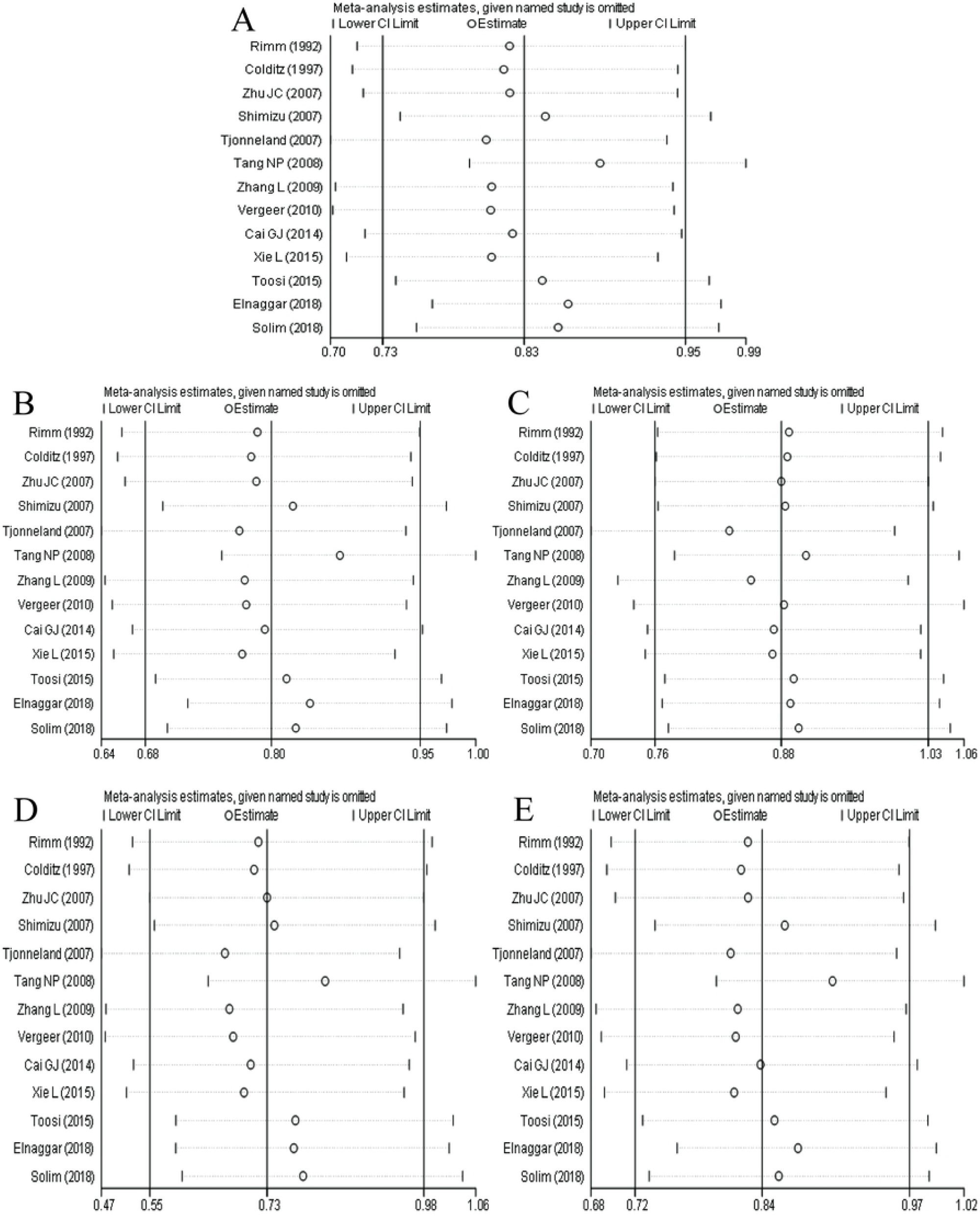
The result of sensitivity analysis between Endothelial lipase 584C/T polymorphism and susceptibility to CAD(A:Allele model; B:dominant model; C:recessive model; D:homozygous model; E:heterozygous model)

## 4 Discussion

The purpose of this study was to explore the association between AGT M235T, EL 584C/T polymorphisms and CAD. In recent years, the polymorphism of AGT and EL genes have attracted considerable attention of researchers. It has been found that both AGT and EL genes have gene polymorphisms, which may affect gene transcription and expression, and are closely associated with the significant increase of coronary heart disease risks [46, 47]. Also, AGT polymorphism may affect restenosis after stent implantation in patients with coronary heart disease [48]. Junusbekov et al. [49] suggests that AGT rs699 gene mutations are associated with cardiovascular phenotypes in atherosclerotic peripheral arterial obstructive disease. Moreover, AGT polymorphism was associated with cardiovascular risk in patients with acromegaly [50]. Studies have shown that serum concentration of full-length endothelial lipase and carboxyl-terminal fragments could predict cardiovascular risk in patients with coronary heart disease [51]. Therefore, it is of considerable significance to study the polymorphism of AGT and EL genes and the susceptibility to CAD.

The results of AGT M235T polymorphism and CAD showed that the differences were statistically significant in the dominant, recessive, homozygous, heterozygous and allelic gene models. Allele T could be a risk factor for CAD. After removing two studies, in which the control group did not satisfy HWE, the results of meta-analysis were consistent with those before exclusion. The Ethnicity subgroup analysis results suggested that except the heterozygous genetic model of the Chinese population, it was statistically significant in other genetic models of the Chinese population and all genetic models of the Caucasian population. Also, the funnel plot of publication bias was symmetrical, and the P-value of Egger’s Test was greater than 0.05, suggesting that there was no publication bias and the correlation between AGT M235T polymorphism and CAD susceptibility was robust, which was also verified by sensitivity analysis results. In a meta-analysis involving nine studies, Wang et al. [52] indicated that the genotype TT of AGT M235T might increase the risk of CAD, which is consistent with the results of this study.

The results of EL 584C/T polymorphism and CAD showed that the differences were statistically significant in dominant, homozygous, heterozygous and allelic gene models, and the recessive genetic model had no statistical significance. Allele model and dominant gene model were still statistically significant after the control groups not meeting HWE were removed, which indicated that the two models were robust. While, in the results of the Ethnicity subgroup analysis, there was no statistical significance in dominant, recessive, homozygous and heterozygous genetic models of Asian and Caucasian populations. Also, the heterogeneity did not decrease significantly, suggesting that Ethnicity may not be the primary source of heterogeneity. The results of publication bias showed that the symmetry of funnel plots in many genetic models were general, and the P values of Egger’s Test were all slightly less than 0.05, indicating the robustness deviation of the conclusion. Also, there may be a potential publication bias having a significant impact on the outcome. It can also be seen from the results of sensitivity analysis that, except the allele genetic model, the differences in other genetic models were statistically significant after excluding some of the literatures, which indicated that the conclusion of 584C/T polymorphism of the EL gene and susceptibility to CAD should be taken seriously. A meta-analysis of EL 584C/T polymorphism and coronary heart disease susceptibility by Cai et al. [53] suggested that there were significant differences between allele model and homozygote model in the Asian population. At the same time, there were no statistical differences in allele model of the Caucasian population, which is not consistent with the conclusion of this study, as more high-quality researches are included in this study.

Furthermore, this study also had some limitations, 1) there was heterogeneity in allele model, recessive gene model and homozygous model of AGT M235T gene, and the heterogeneity did not decrease significantly according to ethnic subgroup analysis, 2) the heterogeneity of the EL 584C/T gene was significant in all genetic models except the recessive gene model, and there was no significant decrease in the heterogeneity after analysis by ethnic subgroup, 3) except recessive gene model, EL 584C/T gene had publication bias in other genetic models, especially in Caucasian population, 4) the interaction between gene and gene, gene and environment was not analyzed.

In conclusion, AGT M235T and EL 584C/T genes are associated with CAD susceptibility. Allele T, genotype TT and TC of AGT M235T gene and allele C, genotype CC of EL 584C/T gene could increase the risk of CAD. In particular, the relationship between the AGT M235T gene and CAD is robust, while the relationship between the EL 584C/T gene and CAD susceptibility is unstable. Furthermore, considering the limitations of this study, further studies are required on a larger scale to explore the relationship between AGT and EL polymorphisms, and CAD susceptibility.

## Acknowledgments

None.

